# altAFplotter: a web app for reliable UPD detection in NGS diagnostics

**DOI:** 10.1101/2023.08.08.546838

**Authors:** Maximilian Radtke, Johanna Moch, Julia Hentschel, Isabell Schumann

## Abstract

The detection of uniparental disomies (the inheritance of both chromosome homologues from a single parent, UPDs) is not part of most standard or commercial NGS-pipelines in human genetics and thus a common gap in NGS diagnostics. To address this we developed a tool for UPD-detection based on panel, exome or whole genome sequencing data which is easy to use and publicly available. Detection of UPDs and delineation of different UPD types is achieved by combining two approaches: detection of runs of homozygosity and their extent per chromosome and investigation of variant inheritance and the ratios of uniparentally inherited variants per chromosome. Both metrics have been carefully tuned with a large set of positive and negative controls to allow reliable and sensitive detection of UPDs. We provide a web tool that can be easily accessed and used to enable geneticists to perform a sensitive UPD-detection analysis based on exome, panel and whole genome sequencing data (.vcf-files) on the fly.

## Motivation

Despite the rapid advancements in next generation sequencing techniques and continously reducing costs, the lack of bioinformatic expertise in smaller labs prevents the application of certain diagnostic methods. These include the detection of uniparental disomies which are of high clinical relevance^1^ and can be identified in large panel, exome or genome sequencing.

In order to enable clinicians and researchers to perform such additional diagnostic steps, we aim to provide a curated application that is easy to use and provides a guideline for interpretation of complex genotypes. The web app „altAFplotter” (https://altafplotter.uni-leipzig.de/) performs a set of analyses and summarizes the results in an overview table but also allows detailed visual exploration of alternative allele frequencies, inheritance patterns and runs of homozygosity (ROH) in an interactive user interface.The app is designed in a way that it can be easily hosted locally and integrated in an existing workflow, as it processes standard vcf files.

## Methods

The identification of UPDs and their classification as isodisomy, heterodisomy, mixed or segmental iso-and heterodisomy can be achieved by examination of ROHs and inheritance patterns per chromosome. For this purpose, we utilize bcftools isec^2^ and bcftools roh^3^. A batch evaluation positive controls from previously described cases^4^ and our patient cohort of ca. 9000 large panel and exome sequencing samples (manuscript in preparation) was used to determine cutoffs for chromosome flagging.

The cutoffs for flagging were selected to ensure highly sensitive detection (27/27 positive controls are detected, see **Error! Reference source not found**., A-C) at the cost of increased false positives (2% in our exome cohort analysis, excluding consanguinous individuals). As the tool is designed for case by case evaluation with manual inspection, we reason that a high sensitivity is the appropriate approach for a diagnostic setting.

Chromosome flagging is informed by the applied method, ROH detection and inheritance ratio:

- Runs of homozygosity: Flags are applied, if the chromosome is covered by >70% (**roh_high**) or between 20%-70% (**roh_mixed**) ROHs. Applicable for both, single and multi-sample analyses.
- Inheritance ratio: This value describes the ratio between maternal and paternal variants and vice versa in trio setups. For duos (index and one parent) it describes the ratio between maternal and not-maternal or paternal and not-paternal variants. Cutoffs here were chosen as follows: >2 for trios and >5 for duos. For these chromosomes the flag **inh_ratio_high** is applied. This allows the detection of iso- and heterodisomies and the identification of the parental origin.
- Consanguinity: if more than three chromosomes exceed a ROH coverage of 10% per patient, the flag **consanguinity_likely** is applied. Such cases can not be reliably evaluated by this approach.
- Insufficient SNVs: If per chromosome less than 200 SNVs are present, the chromosome is excluded from analysis and the flag **insufficient_snvs** is applied.

For interpretation of chromosome flags and their various combinations, table 1 can be consulted. Large deletions also lead to longer ROHs, therefore an additional hint is given to the user to check for those, if a ROH-flag is applied. For this reason and as a general recommendation, validation of UPDs by a second method is strongly advised. Besides the flagging, detailed and interactive plots are available to allow for investigation of affected chromosomes (see Figure S1).

**Table 1:** Interpretation guidelines for chromosome specific UPD-flags as shown in the web-app.

|  |  |
| --- | --- |
| roh_high | potential isodisomy - check for deletions |
| roh_high_mixed | potential isodisomy or mixed UPD - check for deletions |
| inh_ratio_high | potential heterodisomy |
| roh_high(_mixed) + inh_ratio_high | potential isodisomy or mixed UPD |

## Availability and implementation

The altAFplotter-web app is written in python, using streamlit (https://github.com/streamlit/streamlit) as an web-app-framework. It utilizes bcftools via subprocesses to calculate ROHs and variant intersections between vcf files.

All uploaded vcf-files are deleted as soon as the calculations have been performed.

The app is freely available under https://github.com/maxmilianr/altafplotter_public and publicly hosted under https://altafplotter.uni-leipzig.de/.

As altAFplotter is part of our internal diagnostic pipeline, we plan for regular updates and addition of new features. At the time of writing, next upcoming changes include: support for ROH calling with a public variant reference (e.g. gnomAD) for more relevant ROH detection; additional visual clues for UPD interpretation, such as highlighting known clinically relevant UPD regions; improve detection of consanguinity; increased sensitivity for duo setups, where the parent of whom the UPD arised is not sequenced; update to current streamlit version; better integration of whole genome data.

User guidelines for general handling and interpretation of the results can be found here: https://github.com/maxmilianr/altafplotter_public/blob/main/user_guideline/user_guideline.md

## Acknowledgement

We are thankful to Christian Gilissen, Thomas Eggermann and Thomas Liehr for sharing data.

## Author contribution

MR and IS conceived the initial study concept. JM collected controls and determined cut off values; MR wrote and edited the script of the altAF-plotter. MR wrote the initial draft of the manuscript; all authors reviewed and commented on the final draft of the manuscript.

**Figure 1:**
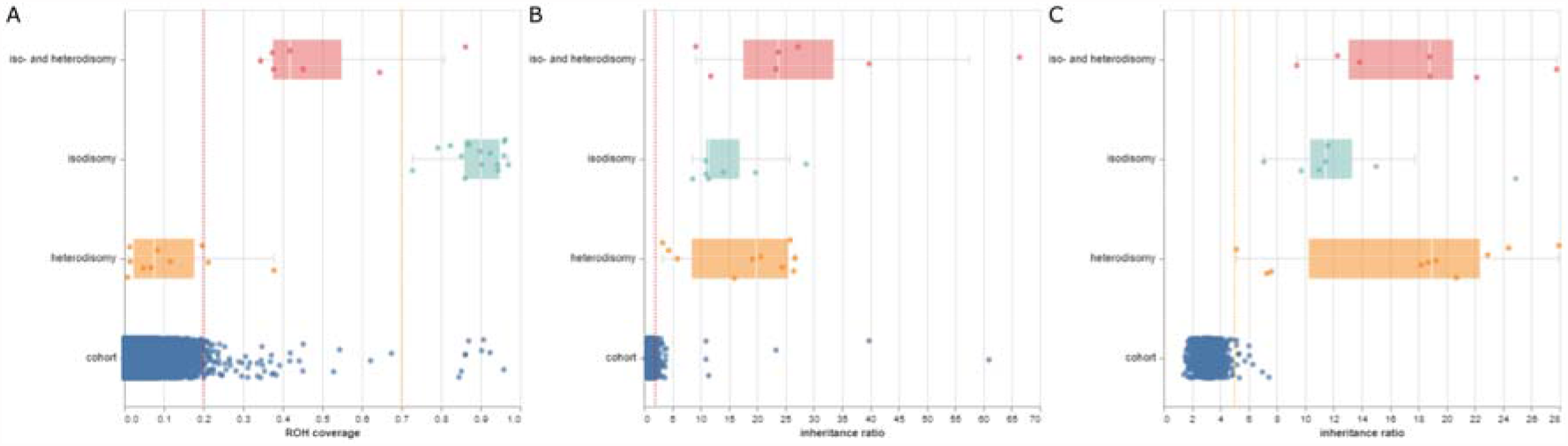
Determination of cutoffs, each data point represents the respective metric (ROH coverage or inheritance ratio) on one chromosome. „Cohort” refers to an evaluation of ∼6200 whole exomes (a subset of the entire cohort: only whole exomes, no consanguinous patients), „iso- and heterodisomy”, „isodisomy” and „heterodisomy” refers to positive controls used to defined cutoffs. (A) ROH coverage cutoffs as defined for mixed UPDs (0.7, orange line) and isodisomies (0.2, red line). Heterodisomies can not be identified based on ROHs. (B) inheritance ratios for trio analyses, shown here is the ratio of maternal over paternal variants. The cutoff (2, red line) includes all positive controls and can be used to identify all three types of UPDs. (C) For duo-analyses, the ratio of maternal over non-maternal or paternal over non-paternal variants can be used to identify UPDs. The cutoff (5, orange line) is chosen to include all positive controls.

