## Supplemental Figure 1 for "altAFplotter: a web app for reliable UPD detection in NGS diagnostics"

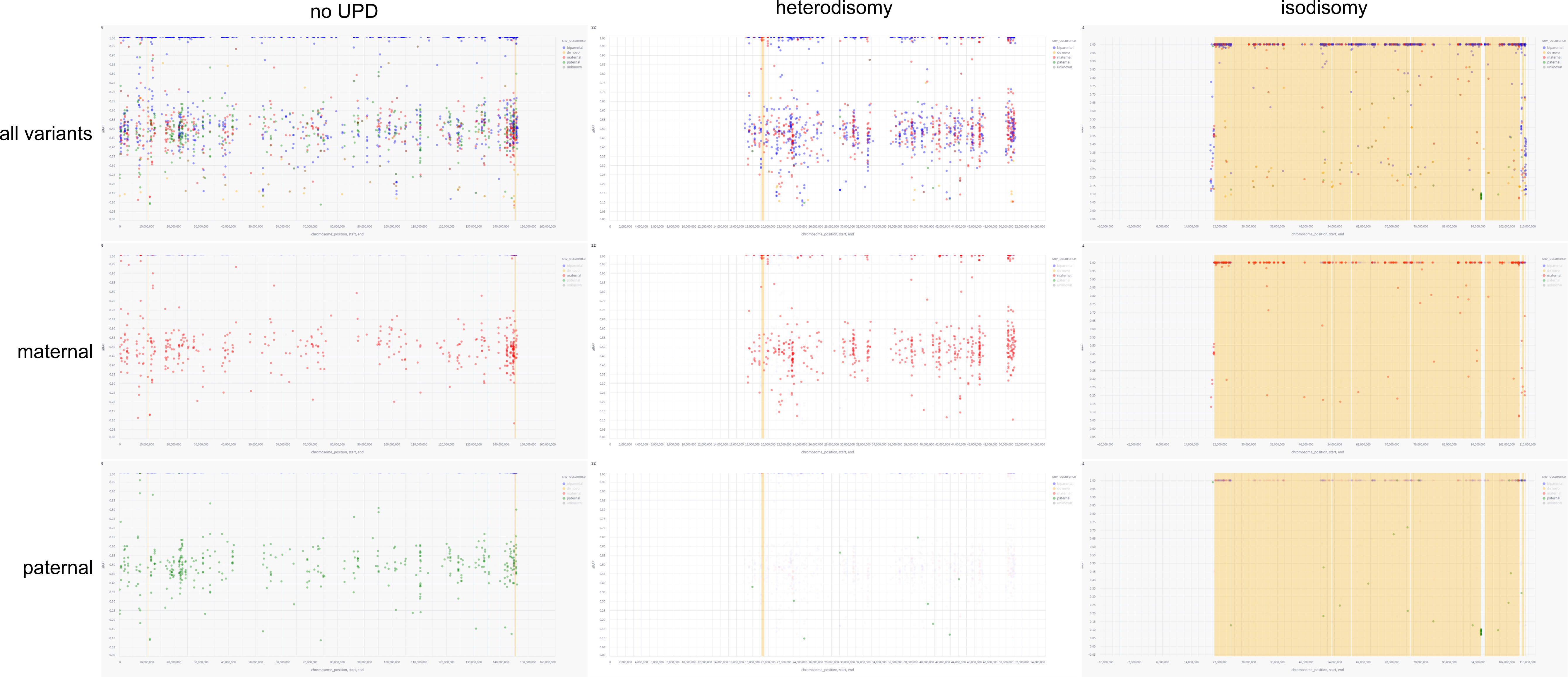

Figure S1: Screenshots of the chromosome plots as shown in the web app. The columns represent three different chromosomes from three different samples. „no UPD“ refers to an unaffected chromosome 8; „heterodisomy“ shows a heterodisomic chromosome 22; „isodisomy“ shows a isodisomic chromosome 4. The rows refer to a different selection of variants to be displayed. The legend refers to the possible variant origins: blue = biparental, yellow = de novo, red = maternal, green = paternal, grey = unknown. The top row „all variants“ shows the allele frequencies of all variants. „Maternal“ and „paternal“ show only those variants that certainly derive from the maternal or paternal allele, respectively. In this view, the origin of the heterodisomy and isodisomy can be easily determined, as only maternal variants can be found on the disomic chromosomes. The orange blocks on the isodisomic chromosome indicate ROHs, which are not present on the heterodisomic chromosome. This stresses the fact that this type of disomy can only be identified in a duo or trio analysis.
